# Confidence of probabilistic predictions modulates the cortical response to pain

**DOI:** 10.1101/2022.08.11.503296

**Authors:** Dounia Mulders, Ben Seymour, André Mouraux, Flavia Mancini

## Abstract

Pain typically evolves over time and the brain needs to learn this temporal evolution to predict how pain is likely to change in the future and orient behavior. This process is termed temporal statistical learning (TSL). Recently, it has been shown that TSL for pain sequences can be achieved using optimal Bayesian inference, which is encoded in somatosensory processing regions. Here, we investigate whether the confidence of these probabilistic predictions modulates the EEG response to noxious stimuli, using a TSL task. Confidence measures the uncertainty about the probabilistic prediction, irrespective of its actual outcome. Bayesian models dictate that the confidence about probabilistic predictions should be integrated with incoming inputs and weight learning, such that it modulates the early components of the EEG responses to noxious stimuli, and this should be captured by a negative correlation: when confidence is higher, the early neural responses are smaller as the brain relies more on expectations/predictions and less on sensory inputs (and vice versa). We show that participants were able to predict the sequence transition probabilities using Bayesian inference, with some forgetting. Then, we find that the confidence of these probabilistic predictions was negatively associated with the amplitude of the N2 and P2 components of the Vertex Potential: the more confident were participants about their predictions, the smaller was the Vertex Potential. These results confirm key predictions of a Bayesian learning model and clarify the functional significance of the early EEG responses to nociceptive stimuli, as being implicated in confidence-weighted statistical learning.

**SIGNIFICANCE:** The functional significance of EEG responses to pain has long been debated because of their dramatic variability. This study indicates that such variability can be partly related to the confidence of probabilistic predictions emerging from sequences of pain inputs. The confidence of pain predictions is negatively associated with the cortical EEG responses to pain. This indicates that the brain relies less on sensory inputs when confidence is higher and shows us that confidence-weighted statistical learning modulates the cortical response to pain.

## INTRODUCTION

In order to survive, animals need to minimise their risk of harm and can do so by learning to predict pain and other body threats. Learning to predict threats is necessary to orient behaviour. How does the brain learn to predict pain and aversive states? The majority of previous work has focused on associative learning to predict pain outcomes based on non-pain cues (Atlas et al., 2010;Atlas and Wager, 2012;Jepma et al., 2018;Strube et al., 2021). Associative learning well describes the prediction of isolated, transient threatening events, but is insufficient to characterise learning to predict long-lasting sequences of pain inputs (Mancini et al., 2022), which typically occur in pain conditions (Schulz et al., 2015). When experiencing temporally-evolving pain, the brain needs to learn to predict forthcoming pain based on its past history. Recently, we have shown that learning to predict pain sequences can be achieved using optimal Bayesian inference, in absence of non-pain cues (Mancini et al., 2022). Probabilistic predictions of the frequency of getting pain are encoded in the human primary and secondary cortex, motor cortex and right caudate, whereas their precision is encoded in the right superior parietal cortex.

Bayesian inference frameworks make testable hypotheses about the role of confidence in learning and its effect on neural activity. The confidence and error of neural predictions are dissociable measures of uncertainty. Confidence is a measure of the variability of the prediction, irrespective of the outcome of the prediction. In contrast, the prediction error refers to the discrepancy between a prediction and reality. A Bayesian inference account predicts that the confidence of a probabilistic inference (1) weights learning, (2) is integrated with sensory information at early stages of information processing, and (3) is inversely related with sensory cortical responses (i.e. high confidence reduces sensory responses) as the brain relies less on incoming sensory inputs (Büchel et al., 2014;Seymour and Mancini, 2020). Here, we test these predictions using a TSL task with thermal stimuli and EEG in healthy, human participants.

We focus on the largest wave that can be recorded from EEG in response to transient sensory stimuli: the Vertex Potential (VP) (Cruccu et al., 2008). The VP is typically composed by a biphasic, negative (N2 component) and positive (P2 component) waveform with a characteristic, symmetric scalp distribution with peak over the vertex (Cz-FCz). The VP can be observed for stimuli in virtually any sensory modality (Mouraux and Iannetti, 2009), but despite its ubiquity there is no consensus over its functional significance.

The traditional interpretation is that the VP reflects the intensity of a sensory stimulus (Chen et al.,2001;Cruccu et al., 2008;De Keyser et al., 2018). A recent study using a pain conditioning paradigm did not find evidence for a modulation of the VP by expectations and prediction errors, suggesting that the VP mostly reflects the sensory processing of a stimulus (Nickel et al., 2022). However, other studies have shown that the amplitude of the VP is modulated by the history and unpredictability of previous stimuli, and can be decoupled from perceived intensity (Bromm and Treede, 1987;Ronga et al., 2012;Torta et al., 2012;Valentini et al., 2012;Mancini et al., 2018).

The seemingly divergent conclusions of previous studies could stem from the different definitions of stimulus predictability and uncertainty, and the lack of a mathematical quantification of these concepts. Here we use a normative approach to dissect the contributions of temporal predictions, their confidence and error on the Event Related Potentials (ERPs) elicited by sequences of somatosensory, thermal stimuli. The stimulus sequences had a probabilistic (Markovian) temporal structure, with underlying statistics that can be learned (Fig.1) (Mancini et al., 2022).

**Figure 1.**
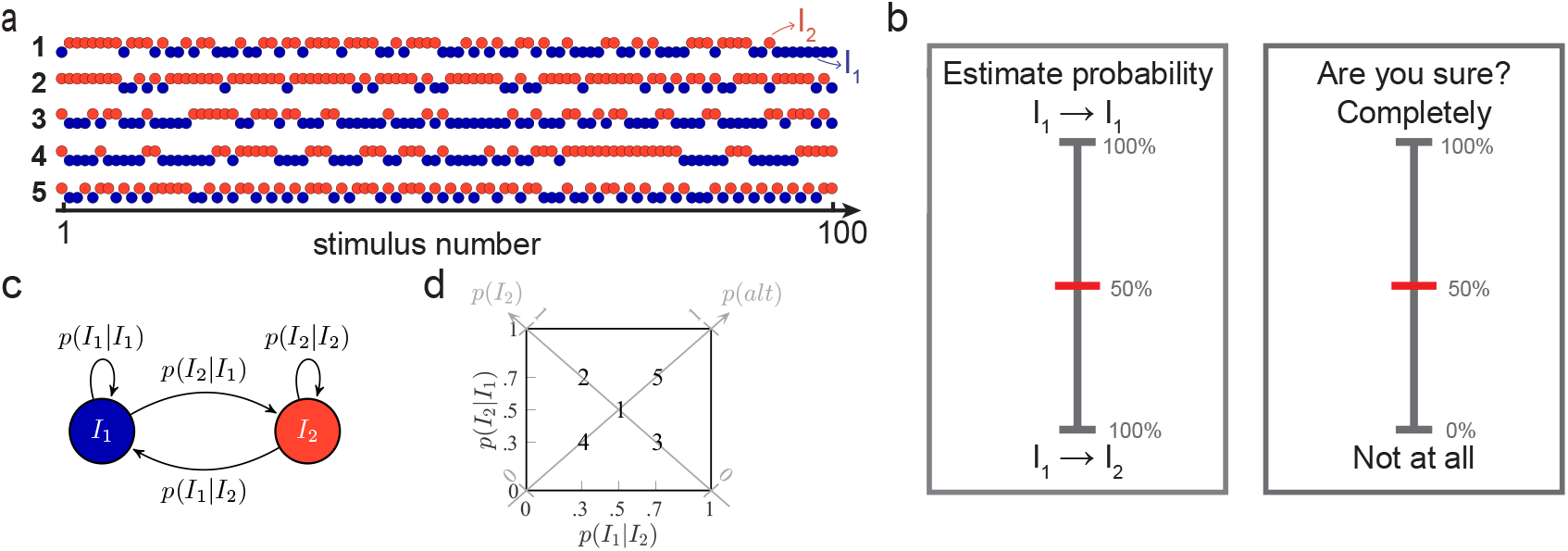
Temporal statistical learning experiment. **a**, Examples of sequences of stimuli of intensities *I*_1_ and *I*_2_ that are applied to the participants’ forearm. Each sequence has different generative statistics (a majority of *I*_2_ or *I*_1_, more alternations or repetitions, etc) and the inter-stimulus-interval (ISI) is set to 3 seconds. **b**, Behavioral questions asked to the participants every 15±3 stimuli in the sequences to evaluate their stimulus probability estimates and numerical confidence in these predictions. The sequences are paused during a maximum of 8 seconds per question. **c**, Markovian generative process of the sequences of stimuli whose intensities are *I*_1_ and *I*_2_. **d**, Transition probability matrix in which the five generative pairs of transition probabilities (TPs) employed are indicated with bold numbers. One example of sequence generated with each of these five TPs is shown in **a**.

## RESULTS

Thirty-one human participants received five different types of probabilistic sequences of thermal stimuli delivered with a contact thermode to the right forearm (Fig.1a). In each sequence, there were two types of stimuli – one stimulus was cold (*I*_1_), and the other was painfully hot (*I*_2_, above the Aδ-fiber threshold). The low intensity was chosen as being cold to ensure that the participants were able to discriminate both intensities based on pilot experiments. The sequences transitioned between the cold and hot stimuli according to a Markovian process described with two generative transition probabilities (TPs, Fig.1c-d). Occasionally, the sequence was paused and participants were asked to predict the probability of the next stimulus based on the previous stimuli and to report their confidence in these estimates on a numerical rating scale (Fig.1b). Each participant received 2 sequences of 100 stimuli generated with each of the 5 distinct TPs indicated in Fig.1d in a randomized order and was informed that the sequence statistics changed (see Methods). On average along the whole experiment, participants received similar numbers of stimuli from both intensities and rated similar numbers of transitions from both intensities (Fig.S1). In line with our previous work, participants were able to predict the frequency of the stimulus intensities, as shown by the positive association between generative and rated item frequencies in Fig.2a. Likewise, with a slightly improved accuracy, participants were able to estimate the transition probabilities from one intensity to the other, as indicated in Figs.2b-c. Finally, the subjective confidence reports were quadratically related to the probability estimates: confidence tended to increase for more extreme probability estimates, as previously reported for auditory and visual sequences (Meyniel et al., 2016).

**Figure 2.**
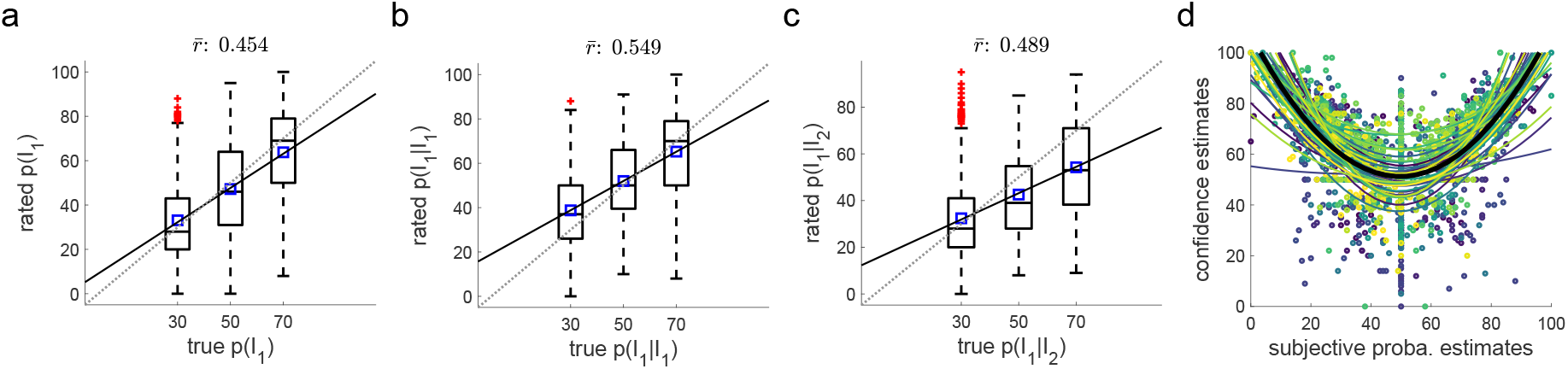
Participants identify the generative sequence statistics. **a**, True and rated probabilities to receive a stimulus of intensity *I*_1_ are correlated subject-wise. The mean correlation across participants is 0.454 (*t*_30_ = 13.603, *p* < 10^−5^, Cohen’s *d* = 2.443), indicating that participants identify the trends within the sequences. Dotted line: identity, plain line: linear fit averaged across participants, blue squares: mean rated probabilities. **b**, Participants also accurately identify the trends in the transitions from *I*_1_. The grand mean correlation between generative and estimated *p*(*I*_1_|*I*_1_) is 0.549 (*t*_30_ = 14.007, *p* < 10^−5^, Cohen’s *d* = 2.516). **c**, Similar to **b** for the transitions from *I*_2_. The grand mean correlation between generative and estimated *p*(*I*_1_|*I*_2_) is 0.489 (*t*_30_ = 11.585, *p* < 10^−5^, Cohen’s *d* = 2.443). **d**, Confidence reports are quadratically related to the probability estimates (mean coefficient of determination of the quadratic fits: *R*^2^ = 0.47). Plain colored lines: individual quadratic fits, thick plain black line: quadratic fit averaged across participants.

### Behavioral modeling

First, we defined the computational principles underlying the participants’ inference of the sequence statistics. We therefore consider a series of models which are fed with the exact same sequences of binary inputs as the participants. Each of these models constructs predictions about the stimulus probabilities along the sequences and can be compared to the subjective reports to shed light on the mechanisms of pain inference.

We fitted two families of three models to the subjective probability estimates obtained in the statistical learning task. One family of models uses Bayesian inference, whereas the other family uses a heuristic, i.e. a non-probabilistic delta rule (Rescorla-Wagner model) with fixed learning rate. The Bayesian models use the confidence of the prediction to weight the update of the representation of the stimulus statistics, whereas delta rule models use a fixed learning rate which is not scaled by uncertainty. In each family, the models differ according to what they predict: the item frequency (IF), the alternation frequency (AF) or the transition probabilities (TPs) of the stimuli.

At group level, we found that probability estimates were best approximated by a Bayesian model which estimates the transition probabilities (Fig 3a). Given that the sequences were not volatile, we used Bayesian models with fixed update of beliefs and a leaky integration to account for forgetting. We estimated that an integration time constant of approximately 8 stimuli best approximated behaviour (Fig 3b), which corresponds to 24 seconds and an integration half-life of around 6 stimuli. This provides evidence that statistical learning for nociceptive stimuli uses a Bayesian inference strategy, whereby the update of the representation is weighted by confidence.

**Figure 3.**
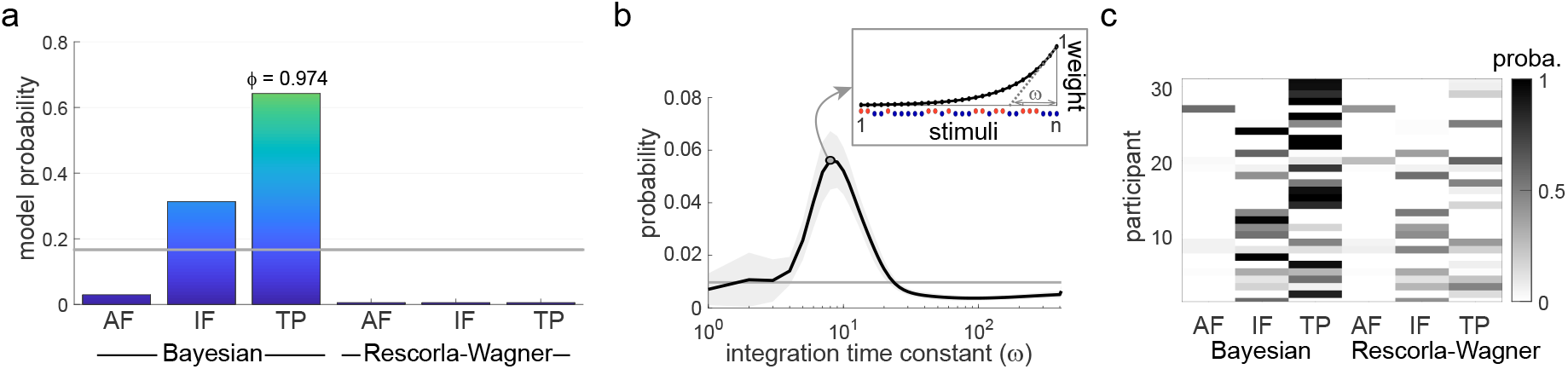
Model comparison. Six different models are considered to explain the subjective reports: Bayesian learners inferring the alternation frequency (AF), the item frequency (IF) or the transition probabilities (TPs), and delta-rule, or Rescorla-Wagner (RW) models, inferring the same sequence statistics (AF, IF, TP). **a**, Bayesian model comparison shows that the participants’ reports are best approximated by a Bayesian model learning the TPs (the exceedance probability of this model – i.e. the probability for this model to be more frequent than the others in the population – is *φ* = 0.974). Colored bars: model probabilities, horizontal gray line: prior (uniform) probability. **b**, Bayesian model averaging reveals that the participants’ integration of observations is best approximated with a time constant *ω* of 8 stimuli. Horizontal line: uniform prior probability, shaded area: s.e.m. across participants, plain dot: curve maximum. The inset illustrates the exponentially decreasing weights that are used to count the number of past stimuli when *n* stimuli have been delivered, with a time constant *ω* of 8. **c**, Individual model probabilities (reflecting the similarity between estimated and modeled probabilities) indicate that most subjective reports are best approximated by the Bayesian model learning the TPs, and to a lesser extent by the Bayesian model learning the IFs, but not much by RW models.

A minority of subjects (n = 11) favoured a simpler Bayesian inference strategy, predicting item frequencies instead of transition probabilities (Fig. 3c). This somehow contrasts with our previous study with volatile sequences, in which only a minority of participants could predict the TPs between the stimuli, whereas the majority of participants showed a preference for the simpler strategy of encoding the IF (Mancini et al., 2022). Here, the two models that best approximate the subjective reports and are above the prior uniform probability remain the Bayesian models learning the IF or the TPs, but most participants were able to predict the more complex temporal statistics that are the TPs (Fig. 3c). This discrepancy can be explained by the fact that the present task was simplified by the absence of volatility in the generative sequence statistics. Note that frequency can always be derived from transition probabilities (the IF corresponds to the principal diagonal of the TP matrix, see Fig. 1d), so participants who prefer a transition probability inference strategy should also access the frequency of the stimuli.

To explore the quality of the fit (i.e. to which extent the winning model is actually close to the participant’s responses), we display the positive correlation between rated and model probability estimates in Fig. 4a. Overall, participants’ reports were highly correlated with the model estimates (grand mean correlation of 0.659, *t*_30_ = 24.4, *p* < 10^−5^). Importantly, the confidence ratings (which were not used to optimize the fit of the model) correlated with the confidence measures deduced from the Bayesian model, Fig. 4b (grand mean correlation of 0.285, *t*_30_ = 9.3, *p* < 10^−5^). Bayesian confidence relates to the statistical certainty about the estimated TPs, i.e. to the inverse spread of the posterior distribution over these TPs. The quality of the confidence fit was similar to previous works (Meyniel, 2020). We then quantified the accuracy of probability and confidence ratings as the correlation coefficients between rated and model estimates, and found they were positively correlated across participants (Fig. 4c, correlation of 0.493, *p* = 0.005). This indicates that optimizing the model to probability estimates provides a good description of participant’s confidence ratings; it also suggests that confidence and probability estimates are derived from a common cognitive process, in line with previous works (Meyniel et al., 2015;Gherman and Philiastides, 2015). Finally, Fig. 4d illustrates the quadratic relationship between Bayesian model probability estimates and confidence, similarly to what we observed for the subjective reports (Fig. 2d).

**Figure 4.**
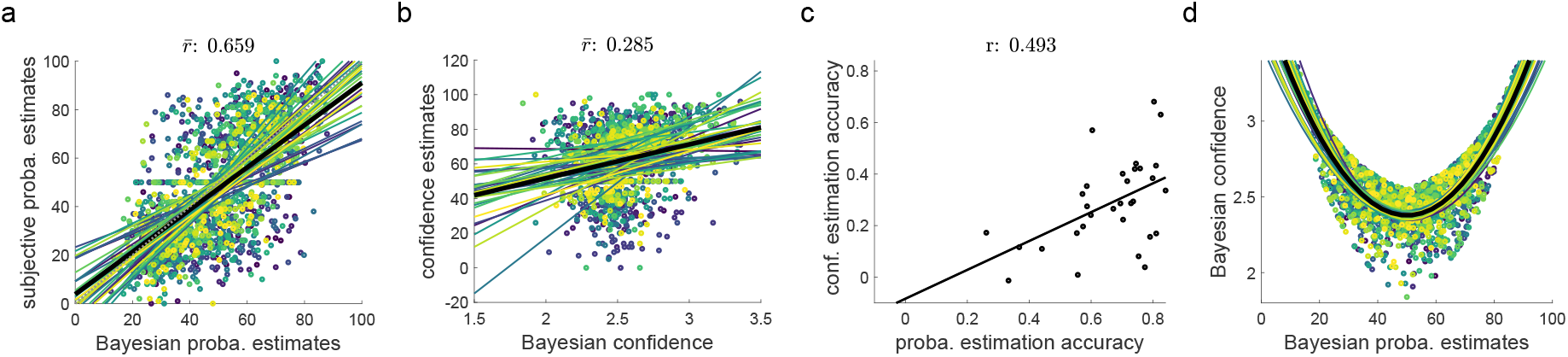
Quality of fit of the best model for the ratings. Subjective estimates of stimulus probability and confidence are highly correlated with Bayes-optimal values obtained from a model learning the TPs with an integration time constant of 8 stimuli. **a**, Scatter plot of estimated and modeled stimulus probabilities, with one color per participant. The grand mean correlation is 0.659 (*t*_30_ = 24.398, *p* < 10^−5^ Cohen’s *d* = 4.382). Dotted line: identity, plain colored lines: individual linear fits, thick plain black line: linear fit averaged across participants. **b**, Scatter plot of estimated and modeled confidence, with the same color code as in **a**. The grand mean correlation is 0.285 (*t*_30_ = 9.293, *p* < 10^−5^, Cohen’s *d* = 1.669). **c**, The accuracy of probability and confidence estimates are positively correlated across participants (Pearson correlation: 0.493, *p* = 0.005). Each accuracy was computed as the correlation coefficient between the subjective reports and the model estimates across trials. **d**, Bayesian confidence is quadratically related to Bayesian probability estimates (mean coefficient of determination ofthe quadratic fits: *R*^2^= 0.59). Plain colored lines: quadratic fits obtained using the sequences of each participant, thick plain black line: quadratic fit averaged across participants’ sequences.

### EEG

Sixty-four channels EEG was recorded on all participants while they were exposed to the sequences of thermal stimuli. As expected, the main evoked response consisted in a biphasic waveform – the Vertex Potential (VP) – which peaked over fronto-central electrodes (Cruccu et al., 2008;Legrain et al., 2011). Figure 5a illustrates the grand-average VPs following cool (*I*_1_) and hot (*I*_2_) stimuli, with scalp topographies of their two main components: the N2 and P2 waves. These two components peaked at 205 ±17 ms and 318 ±40 ms after stimulus onset for *I*_1_, and 369 ±33 ms and 518 ±42 ms for *I*_2_ (mean ± s.t.d.), similar to previous studies using thermal stimulation (De Keyser et al., 2018;De Schoenmacker et al., 2022). The VPs in response to both types of stimuli were analyzed separately given their different latencies and thermal qualities. At a single trial level, the earlier N1 wave was not clearly identifiable due to its low signal-to-noise ratio.

**Figure 5.**
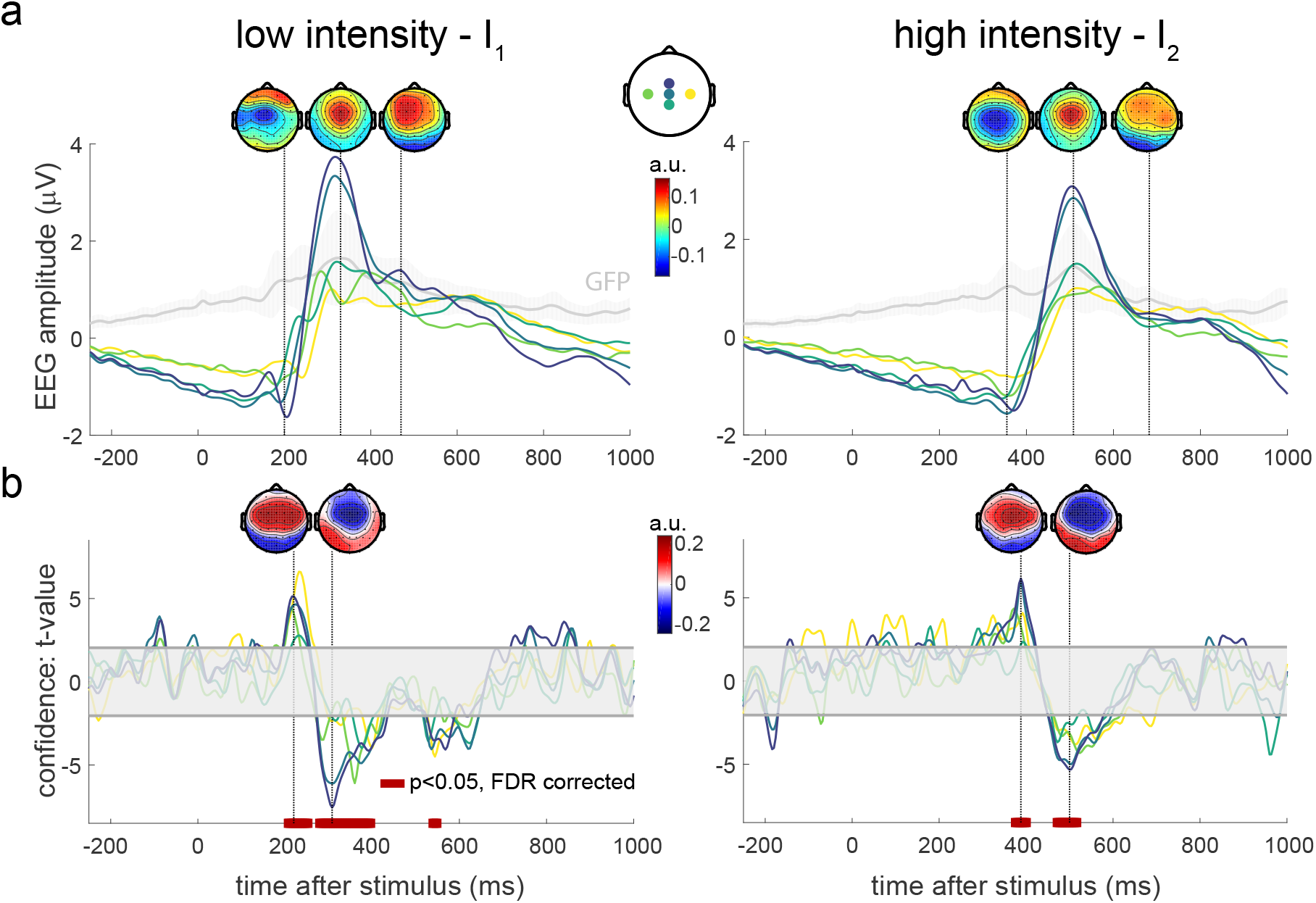
EEG correlates of Bayesian confidence. **a**, EEG responses averaged over trials and blocks, for low (left) and high (right) stimulation intensities. Global Field Power (GFP) time courses are shown in gray, with shaded s.d. across participants. Labels of depicted electrodes: C3, Cz, FCz, CPz, C4. **b**, Encoding of residual confidence in the EEG responses – *t*-statistics for the regression coefficients associated with model confidence. Confidence is obtained from the model which best explains the participants’ behavior: a Bayesian model learning the TPs with an integration time constant of 8 stimuli. The shaded horizontal areas centered around 0 indicate the non-significant regions for *p* < 0.05, two-tailed. Red bars at the bottom of the plots show intervals where the regression coefficients are significantly different from 0 after False Discovery Rate (FDR) correction of the significance levels. Topographies of the largest effects are indicated.

Crucially, we investigated whether the confidence and error of the probabilistic inferences modulate the Vertex Potentials. Using the learning model which best explains the subjective reports (a Bayesian model learning the TPs with an integration time constant of 8 stimuli), we regressed the single-trial EEG signals on two distinct inferential quantities: the residual confidence and Bayesian prediction error (BPE). Confidence is defined as the log precision of the posterior distribution over the latent parameter and is therefore inversely proportional to the posterior variance – confidence gets higher when the variance gets smaller (see (7)). The residual confidence is obtained from the confidence by regressing out the predicted probability, its square and its logarithm to ensure that these quantities do not drive the confidence effects (see Methods and (11)) (Meyniel, 2020). Besides, BPE corresponds to the difference between the received intensity and its predicted probability (see (8)). For each participant, we included these two regressors in linear regressions at each time point from −0.5 to 1 second around stimulus onset and at central electrodes of interest (C3, Cz, FCz, CPz, C4). To make sure that BPE and confidence were not collinear, confidence was regressed on BPE subject-wise, leading to average variance inflation factors (VIFs) of 1 and 1 for *I*_1_ and *I*_2_ respectively, (regression *R*^2^ < 10^−5^). Two variables are typically considered to be highly collinear when their VIF is above 5 (Sheather, 2009).

Grand averages of the *t*-statistics obtained from *t*-tests against 0 for the regression coefficients are shown in Figs. 5b and 6. First, we found a clear modulation of the VP by residual confidence for both intensities (Fig.5b). The sign of these modulations is opposite to the VP, meaning that the larger the confidence, the smaller the N2 and P2 components.

**Figure 6.**
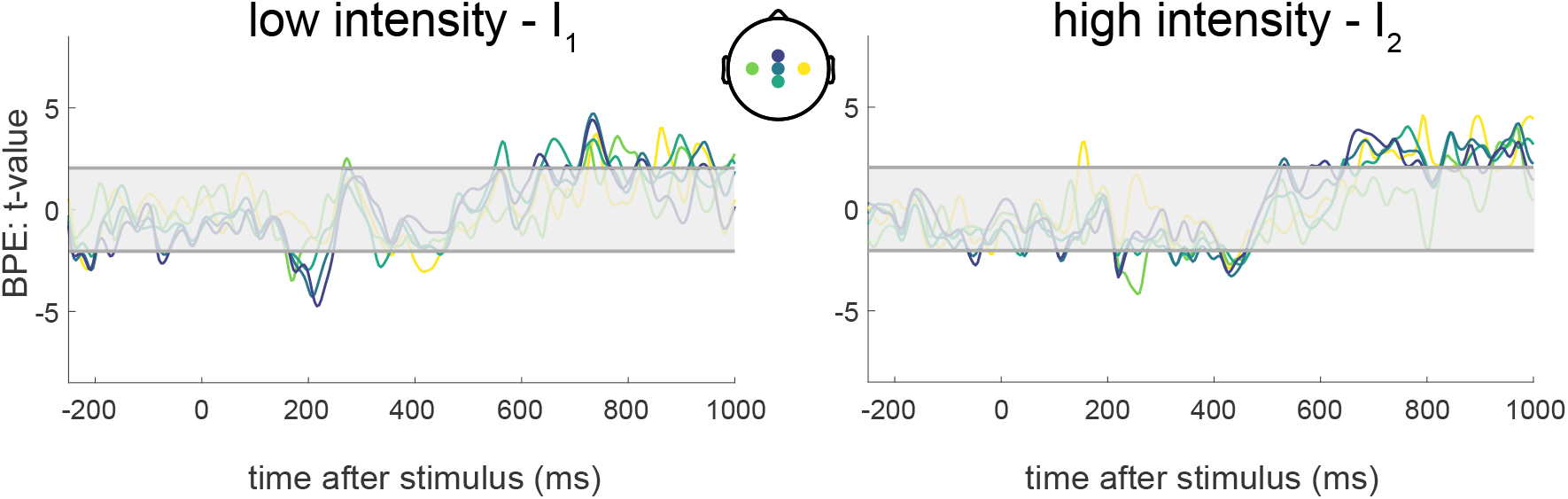
EEG correlates of Bayesian prediction errors (BPE). Encoding of BPE in the EEG responses, similar to Fig. 5b – *t*-statistics for the regression coefficients associated with BPE. BPE is obtained from the model which best explains the participants’ behavior: a Bayesian model learning the TPs with a time constant of 8 stimuli. The shaded horizontal areas centered around 0 indicate the non-significant regions for *p* < 0.05, two-tailed. No time interval was deemed significant after False Discovery Rate (FDR) correction of the significance levels.

Supplementary analyses show that using confidence instead of residual confidence leads to comparable observations (Fig.S2, even though the VIFs are slightly larger in this case). If the Bayesian model learning the IF instead of the TPs is considered (second best model fitted to the behavioral reports), results are also similar (Fig. S3).

Finally, we found no statistical evidence for a modulation of the BPE on the EEG potentials, after correcting for the False Discovery Rate (Fig. 6). However, the prediction error derived from a Bayesian model learning the IF instead of the TPs significantly modulates late EEG waves (Fig. S3). The IF model typically leads to more confident predictions than the TP model, because it is simply inferring one parameter (the frequency) rather than two transition probabilities. However, the IF model predictions are more likely to be ‘wrong’ than the TP model predictions, because the sequences of stimuli were generated using TPs rather than only IFs. Bigger BPEs should yield stronger modulations of the late EEG waves, according to a hierarchical Bayesian inference framework. This is what we find, i.e. the IF BPE modulates more consistently late cortical responses than the TP BPE.

## DISCUSSION

The brain needs to learn to predict forthcoming nociceptive stimuli in order to minimize potential harm. When pain persists over time, the brain needs to extract and learn structure or patterns from streams of sensory inputs without relying on explicit feedback or associated cues (Giorgio et al., 2018). Using a statistical learning task in conjunction with EEG, we provide evidence in support of the view that the human brain uses confidence-weighted Bayesian inference to learn to predict future pain levels and that confidence modulates the cortical response to pain. (Yoshida et al., 2013;Grahl et al., 2018;Valentini et al.,2011;Brown et al., 2008). First, we found that subjective probability estimates of thermal sensations and the associated confidence reports are well approximated by a Bayesian inference model. The best fitting model learns the transition probabilities within the sequences and accounts for participant’s forgetting by integrating past observations with a time constant of 8 stimuli (24 seconds). At the opposite of non-Bayesian models, this winning model indicates that the effect of prior expectations is weighted by confidence to predict forthcoming nociceptive inputs (Meyniel and Dehaene, 2017;Jepma et al., 2018;Mancini et al.,2022). Second, the modeled confidence was negatively associated with the amplitude of the Vertex Potential (VP): the higher the participants’ confidence in the intensity prediction, the smaller the VP. Prediction Errors (PEs), measuring the discrepancy between the expected stimulus and the one which was received, were only weakly associated with increases in later EEG responses. These findings were predicted by our hierarchical Bayesian processing hypothesis: high confidence reduces the cortical response to thermal stimuli because the brain relies less on incoming sensory information, and more on prior information, to generate an inference.

The notion of confidence corresponds to a ‘feeling-of-knowing’ about some variables in an uncertain environment (Meyniel et al., 2015). It is important to note that this notion is employed in two kinds of situations, leading to different computational definitions of confidence. First, confidence in a discrete variable that is learned can be quantified by the probability for this variable to take a given value; it corresponds to the so-called choice or decision confidence (Kepecs et al., 2008;Hangya et al., 2016;Sanders et al., 2016;Herding et al., 2019;Pouget et al., 2016). Second, confidence in the value of a continuous variable instead relates to the spread (often quantified by the standard deviation) of the estimated posterior distribution of this variable (Meyniel et al., 2015;Lebreton et al., 2015;Pouget et al., 2016). For instance, in a TSL task like in this work, the confidence in the next stimulus intensity corresponds to the estimated probability to receive this intensity, while the confidence in the sequence statistic that is learned (AF, IF or TP) is related to its estimated standard deviation. As a consequence, decision confidence – which has been the object of numerous publications about choice and decision-making – should not be confounded with the inferential confidence studied here. For the EEG analysis presented in Fig.5, the estimated probability of each intensity has even been regressed out to obtain the *residual* confidence which is not linearly nor quadratically related to decision confidence.

Statistical models of sensory perception predict that inferential confidence should serve as a weighting factor increasing the effect of prior beliefs on perception (Brown et al., 2008;Büchel et al., 2014; Meyniel and Dehaene, 2017). In the pain field, a few works have studied this principle: from a behavioral view point, confidence indeed modulates pain perception by weighting the effect of expectations (Brown et al., 2008;Grahl et al., 2018;Yoshida et al., 2013). While it is clear that individuals are able to provide metacognitive judgments about pain to some extent (Dildine et al., 2020), some works suggested that humans have a less accurate sense of confidence in the sensory discrimination of pain compared to other sensory modalities (Beck et al., 2019). This contrasts with our finding that inferential confidence is correlated with the Bayesian model confidence, suggesting it is derived from a near-optimal inference process.

Regarding the effects of confidence on brain response dynamics, in a hierarchical Bayesian framework we would expect to see early modulations of EEG responses by confidence, such that increased confidence would lead to a reduction of these responses (Brown et al., 2008;Seymour and Mancini, 2020). The few existing studies that looked at confidence effects on EEG signals are consistent with this view (Valentini et al.,2011;Brown et al., 2008), but haven’t tested its key predictions on the main EEG responses to pain. Here, we show that confidence in statistical inference has a negative association with an early cortical response to nociceptive stimuli, i.e. the VP. The functional significance of the VP has been debated for decades. Traditionally, it was thought that the VP reflects the sensory processing of a stimulus, and it is indeed often used in clinical neurophysiology as a marker of sensory function (Chen et al., 2001;Cruccu et al., 2008;De Keyser et al., 2018). Using nociceptive stimuli, the VP has been associated with subjective pain intensity and, as such, it could be influenced by perceptual and attentional mechanisms (Garcia-Larrea et al., 1997;Lee et al., 2009). Other works have shown that the VP is more likely to encode the differential intensity of a stimulus (with respect to baseline) rather than its absolute intensity (Somervail et al., 2021). Besides, several studies have emphasized that the VP amplitude is not only affected by stimulus intensity and the recent history of stimulation, but also by the unpredictability, novelty and saliency of each stimulus (Iannetti et al., 2008;Valentini et al., 2011;Zhang et al., 2012; Ronga et al., 2012). For instance, just repeating the same stimulus a few times induces a dramatic habituation of the VP, despite the fact that perception remains stable and peripheral habituation can largely be ruled out (e.g. because a new skin spot has been stimulated after each stimulus) (Iannetti et al., 2008;Mancini et al., 2018). Still, a more recent study using a cued pain paradigm suggested that the VP is mostly associated with the sensory processing of a stimulus, without being affected by expectations and PEs (Nickel et al., 2022). These different interpretations can result from the lack of a computational quantification of the pain learning process on a trial basis that would enable fitting individual learning models to each participant (Karlaftis et al., 2019;Beck et al., 2019). Indeed, the aforementioned works did not have estimates of uncertainty or confidence at an individual level because they relied on axiomatic approaches and/or cue-based paradigms. Here, we introduce a computational approach which quantifies nociceptive inference trial-by-trial, enabling the direct correlation of information processing quantities to their brain encoders instead of limiting the contextual information to binary intensities or discrete stimulus and cues categories.

Another component of the statistical learning process is the generation of prediction errors (PEs), measuring the difference between what is predicted (based on previous experiences) and what is actually received. PEs (or surprise) signals are expected to modulate some brain responses regardless of the sensory modality (Maheu et al., 2019), though it is likely that the neural implementation of these effects have some stimulus-specificity (Frost et al., 2015). Here, we did not find significant evidence for an effect of PE on the VP, although there was a weak modulation of late-onset EEG responses. In different paradigms, using shorter sequences of stimuli, PEs can account for shorter time-scale habituation (Somervail et al., 2021;Strube et al., 2021). This is not incompatible with our findings: in short and/or cued sequences, PEs tend to be large and this is likely to lead to a stronger cortical modulation, as dictated by Bayesian inference.

To conclude, we have shown that subjective probability reports about nociceptive intensity are well approximated by a Bayesian model learning the transition probabilities between high and low intensity stimuli. The Bayesian model’s confidence was correlated with the participants’ reported confidence levels. Importantly, inferential confidence was negatively correlated with the VP – the higher the confidence, the smaller the VP. This indicates that the VP is modulated by confidence-weighted statistical learning of sequences of nociceptive inputs and is consistent with the predictions of a hierarchical Bayesian inference framework. Given that some pathological pain conditions have been associated with altered learning and predictive capabilities (Baliki et al., 2010, 2011; Smith et al., 2008; Ploner et al., 2016), future works could assess how confidence representations are modified in these patients, opening the path to promising translational studies.

## METHODS

### Participants

Thirty-six healthy participants (19 females) took part in the experiment, aged 18-30 years, 32 of them being right-handed. The study was approved by the local ethics committee (Comité d’Ethique Hospitalo-Facultaire de l’Université catholique de Louvain, B403201316436). All participants gave written informed consent and received financial compensation. Five participants were not able to distinguish the two stimulus intensities until the end of the experiment, so they were excluded from all the analyses, leaving 31 subjects (16 females).

### Experiments

The task aims to assess temporal statistical learning (TSL) using sequences of nociceptive stimuli of two distinct intensities – *I*_1_ and *I*_2_. The core principle is that as participants are exposed to a such stream of stimuli, they are able to track the sequence statistics to some extent. Indeed, as the sequence goes, one collects evidences of whether the sequence contains more *I*_1_, more *I*_2_, systematically more *I*_1_ following *I*_2_ or *I*_2_, etc. In our experiment, we aim to understand how these learning mechanisms are implemented.

#### Stimuli and generative model

The stimulus intensity *y_n_* ∈ {*I*_1_, *I*_2_} at each time step *n* along a given sequence is uniquely generated according to a two-state Markovian process such that

- 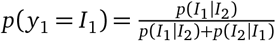
- *p*(*y_n_*|*y*_1:*n*–1_ = *p*(*y_n_*|*y*_*n*–1_

Each sequence is therefore characterized by its generative transition probabilities (TPs, (*p*(*I*_1_|*I*_2_), *p*(*I*_2_|*I*_1_))), i.e. the probabilities of either intensity given the previous stimulus intensity. The stimuli were 250ms-long thermal pulses, applied to the participant’s right volar forearm with a contact thermode (QST Lab, Strasbourg, France, active stimulation surface: 120mm^2^, heating and cooling ramps of 300°/s). To ensure that the participants were able to easily identify the stimulus intensities along all the tested sequences, the low intensity *I*_1_ was chosen to be non painful and cool, while the high intensity *I*_2_ was selected to be painful and above the individual Aδ fiber threshold while being bearable. The temperatures employed were therefore *I*_1_ = 15°C and *I*_2_ = 58°C, up to modifications based on individual thresholds and/or discrimination capabilities, as detailed below. The high intensity *I*_2_ was described as painful and pricking by all participants.

#### Procedure

Each participant underwent the following steps: (1) Aδ fibers threshold estimation through a staircase procedure using reaction times, (2) one pre-check block to assess the discrimination of the two stimulus intensities, (3) one training block, (4) 10 testing blocks and (5) one post-check block to re-assess the discrimination of the two stimulus intensities at the end of the experiment. The total duration of the experiment was approximately 3 hours.

#### Aδ fibers threshold estimation

The threshold for activating Aδ fibers was determined with an adaptive staircase procedure using reaction times (RTs) as described in (Churyukanov et al., 2012). A 250 ms heat stimulus was assumed to activate Aδ fibers when the perception RT was ≤ 650 ms. Starting with a 45°C-stimulus, temperature was increased until the RT became shorter than 650 ms, which led to decrease the next stimulus temperature. The successive absolute temperature differences were in {5, 2, 1, 0.1}°C, decreasing after each detection change (RT shorter vs. longer than 650 ms). The threshold was defined as the mean of 4 stimulation temperatures which led to 3 consecutive changes of RT shorter vs. longer than 650 ms. This led to thresholds of 52.7°C (±5.1) on average (±standard deviation).

#### Check blocks

During each pre-check and post-check block, the participant received a random sequence of 15 stimuli with intensities *I*_1_ and *I*_2_ (fully random TPs of (0.5, 0.5)) and self-paced inter-stimulus-intervals (ISIs). After each stimulus, the participant was asked to report the stimulus identity (cool or hot) and the thermode was displaced before delivering the next stimulus. If there were more than 5 mistakes in a pre-check block, pronounced hesitations about the stimulus identity or if *I*_2_ was unbearable, the stimulus intensities were adjusted accordingly. This led to increase *I*_1_ for 4 participants and decrease *I*_2_ to 57°C for 10 participants. If there were more than 5 mistakes in a post-check block, the subject was excluded from the analyses.

#### Training and testing blocks

During a training or testing block, the participant was exposed to one sequence of stimuli whose intensities were generated based on fixed TPs. The thermode was displaced on the forearm between successive stimuli to avoid trial-to-trial habituation and sensitization which could prevent the participant from distinguishing the two intensities until the sequence ended. The within-sequence ISI was set to 3 seconds. Every 15 ±3 stimuli, the sequence was paused to probe the participant’s inference of the sequence TPs – the participant was asked to (1) estimate the probability of the next stimulus intensity and then (2) rate their confidence in this estimate, Fig. 1b. The scales were displayed on a computer screen in front of the participant and numerical ratings were collected based on keyboard inputs. A time limit of 8 seconds was set to answer each question to avoid too long breaks within the sequences which could affect learning (Atlas et al., 2021).

The **training block** consisted of one sequence of 50 stimuli generated with TPs (0.7, 0.4) and enabled the participants to understand the generative process and familiarize with the task. Subjects received a feedback at the end of this sequence on the correctness of their rating trend.

In each of the **10 testing blocks**, the participant received one sequence of 100 stimuli. The first and last 5 sequences were generated with the 5 different TPs indicated with numbers in Fig. 1d: (0.5, 0.5), (0.3, 0.7), (0.7, 0.3), (0.3, 0.3) and (0.7, 0.7). The order of the blocks was randomized across participants and variable breaks were allowed between sequences.

Behavioral data were analyzed with Matlab R2019b (The MathWorks) and Cohen’s *d* is reported as effect size for each *t*-test.

### Learning models

The generative parameters of the sequence can be continuously estimated based on the stimuli received, leading to predictions about the forthcoming stimulus. To understand how participants perform this inference task, different models performing the same task were fitted to the subjective probability estimates and compared.

Two families of learning models were considered to explain the sequence statistics inference: a Bayesian learner and a non-Bayesian Reinforcement Learning (RL) model which is called the delta-rule or Rescorla-Wagner (RW) model (Meyniel et al., 2016; Meyniel and Dehaene, 2017;Rescorla and Wagner, 1972).

#### Bayesian model

A Bayesian model estimates the posterior distribution of a latent parameter *θ* given the sequence of observed stimuli *y*_1:*n*_ at each time step *n* using Bayes’ rule (Meyniel et al., 2016). Each model *M* estimates specific sequence parameters: either the item frequency (IF) or the alternation frequency (AF) or the transition probabilities (TPs). Given a model *M*, the parameter posterior is obtained by combining the parameter prior and the likelihood of past observations:

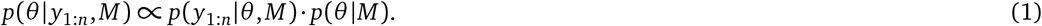

We use a uniform (conjugate) prior distribution over the parameter values (i.e. (*p*(*θ*|*M*) ~ Beta(*θ*|1, 1), which enables deriving analytical solutions for the posterior. Using the Markovian assumption *p*(*y*_*n*+1_|*y*_1:*n*_, *θ*) = *p*(*y*_*n*+1_|*y_n_,θ*), the likelihood can be decomposed as

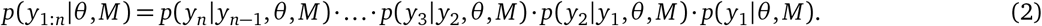

This likelihood and thereby the posterior can be further simplified depending on the model *M* as shown below.

1. **IF learning**. With this model, the inferred parameter is the probability to receive a stimulus of intensity *I*_1_: *θ* = *p*(*I*_1_) := *θ*_*I*_1__. The posterior is therefore

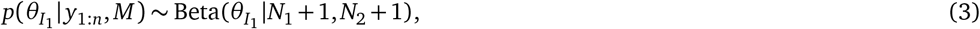

where *N*_1_ and *N*_2_ are the numbers of stimuli of intensity *I*_1_ and *I*_2_ respectively within *y*_1:*n*_.
2. **AF learning**. The inferred parameter is the probability of intensity alternation, i.e. the probability to switch from *I*_1_ to *I*_2_ or vice versa within the sequence: *θ* = *p* (alt.) := *θ*_alt_. The posterior dsitribution reads

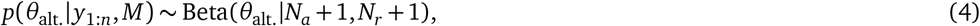

with *N_a_* and *N_r_* the number of alternations and repetitions of stimulus intensities within *y*_1:*n*_.
3. **TPs learning**. The inferred parameter is now two-dimensional and corresponds to the transition probabilities of the sequence of stimuli: *θ* := (*θ*_*I*_1__|_*I*2_, *θ*_*I*2_|*I*_1_), which leads to the posterior

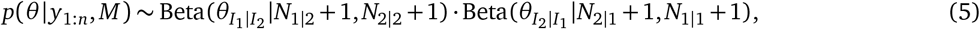

where *N*_*j*|*k*_ is the number of transitions from *I_j_* to *I_k_* counted within *y*_1:*n*_.

To account for limited memory constraints during inference and an unknown timescale of integration, a leaky integration of observations is considered (Meyniel et al., 2016). All the models are endowed with a free parameter *ω* ∈ [1, ∞[– the integration time constant – and the *k*^th^ last observation counted (being it an item, an alternation or a transition depending on the model considered) is weighted according to an exponential decay by a factor exp^−*k*/*ω*^.

For all Bayesian models, some outcomes of interest can be deduced from the posterior at each position *n* within the sequence, when the observations *y*_1:*n*_ have been received:

- **The probability of the next stimulus** is the mean of the posterior distribution:

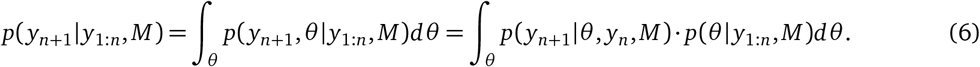
- **The confidence in the learned parameter** relates to the precision (inverse variance, *π* := 1/*σ*^2^) of the posterior (Meyniel and Dehaene, 2017;Pouget et al., 2016):

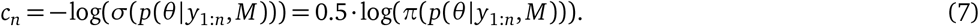 The confidence quantifies the certainty in the estimated continuous variable, and is typically expressed in log space so that the standard deviation and variance are proportional.
- **The prediction error** is defined like in a Bayesian predictive coding framework (Aitchison and Lengyel,2017;Geuter et al., 2017) as

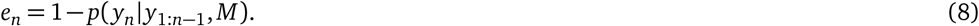 It can be noted that, likewise, the Shannon surprise (Meyniel and Dehaene, 2017) elicited by the last stimulus also quantifies the discrepancy between the intensity that was expected and the one that is received (*y_n_*), in a log space: *S_n_* = –log(*p*(*y_n_*|*y*_1:*n*–1_, *M*)).

To assess the extent to which these models and their parameter (the integration time constant) are identifiable in our experiment, parameter and model recovery analyses can be found in Fig. S4.

#### Rescorla-Wagner, or delta-rule, models

The delta-rule model, or Rescorla-Wagner (RW) model (Rescorla and Wagner, 1972;Miller et al., 1995), is compared to the Bayesian model. While the latter weights the posterior updates by confidence (Meyniel and Dehaene, 2017), the delta rule uses a constant and non-statistical weighting of incoming observations to estimate the latent parameter. The inferred parameter *θ* (IF, AF or TPs) is initiated at 0.5 and is seen as a state value *V* in the RW models, as detailed in what follows.

1. **IF learning**. The state value corresponds to the estimated probability to receive a stimulus of intensity 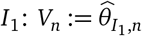. At each step *n* in the sequence, the state is updated as *V_n_* = *V*_*n*–1_ + *α* · (*R_n_* – *V*_*n*–1_), where *R_n_* = 1 if *y_n_* = *I*_1_ and *R_n_* = 0 if *y_n_* = *I*_2_ and with the learning rate *α* ∈]0,1[being a free model parameter.
2. **AF learning**. The state value corresponds to the estimated probability of an alternation within the sequence: 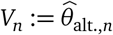. The state is updated as *V_n_* = *V*_*n*–1_ + *α* · (*R_n_* – *V*_*n*–1_), where *R_n_* = 0 if *y_n_* = *y*_*n*–1_ and *R_n_* = 1 otherwise.
3. **TPs learning**. The state value is two-dimensional and corresponds to the estimated transition probabilitiess: 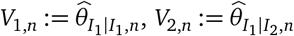.

The state is updated as

- *V_i,n_* = ^V^_*i,n*–1_ + *α* · (*R_n_* – *V*_*i,n*–1_), with *R_n_* = 1 if *y_n_* = *I*_1_ and *R_n_* = 0 if *y_n_* = *I*_2_, if *y_n_* –1 = *I_i_*
- *V*_i,n_ = *V*_*i,n*–1_ if *y*_*n*–1_ = *I_i_*

### Model fitting

To determine to which extent each model accounts for the subjective reports, we quantify the relationship between subjective and model probability estimates by linearly regressing the subjective reports on the modeled estimates for each participant and model. Across trials indexed by *n*, the probability report *x_n_* is hence regressed on the model probability of *I*_1_ 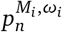 deduced from each model *M_i_* with free parameter *ω_i_* as described above (Bayesian and RW models learning the IF, AF or TPs, with integration time constant or learning rate as free parameter) as:

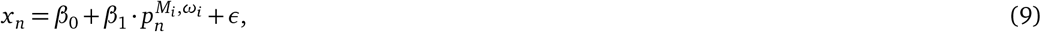

where *β* are the regression coefficients, estimated by OLS, and *ε* the residuals.

The quality of this fit is quantified by the model evidence (or marginal likelihood) *p*(*x*|*M_i_*), which is estimated with the Bayesian Information Criterion (BIC) as:

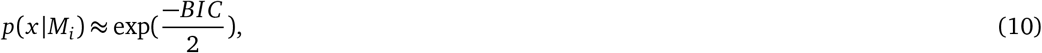

with 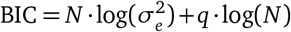, the mean squared error (MSE) of the regression 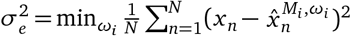, *N* the number of observations and *q* the number of parameters (here there are 2 regression coefficients and 1 model free parameter). When comparing models with the same number of parameters, minimizing the BIC amounts to minimizing the MSE. We considered 99 possible learning rates for the RW models in the range from 0.005 to 0.95, and 103 integration time constants for the Bayesian models from 1 to 400 plus infinity (i.e. a perfect integrator).

Individual, subject-wise, model probabilities were obtained by normalizing the model evidences estimated with the BIC as in (10).

### Model comparison

The model with the largest model evidence (or lowest BIC) was considered to be the best fit for the ratings. To compare the six models *M_i_* described above, we conducted a Bayesian model comparison as implemented in the VBA toolbox (Daunizeau et al., 2014)and adopted a random-effect approach, assuming that the optimal model can differ across participants. The analysis yielded the expected probability of each model *M_i_* and the probability for *M_i_* to be more frequent than all the other models in the population, which is called the ‘exceedance probability’ and is denoted by *φ*.

The model free parameter which approximated the subjective reports best on average was determined through Bayesian model averaging (Maheu et al., 2019) for the Bayesian and RW models separately by estimating *p*(*ω*|*x*) ∝ ∑_i_*p*(*x*|*M_i_*,*ω*) ≈ ∑_*i*_exp(−BIC(*M_i_*,*ω*)/2).

### Electrophysiological recordings

EEG was recorded during the whole experiment using 64 Ag-AgCl electrodes placed on the scalp according to the international 10/10 system (WaveGuard 64-channel cap, Advanced Neuro Technologies) and with an average reference. Synchronization of the stimuli, triggers on the EEG and behavioral questions was performed with the Data Acquisition Toolbox and Psychtoobox running on Matlab. Electrode impedances were kept below 10kΩ. Eye movements were recorded using a pair of surface electrodes placed above and on the right side of the right eye, and one electrocardiogram (EKG) lead was recorded with two surface electrodes, one below the right clavicle near the shoulder and the other on the last left rib. Signals were amplified and digitized at 1000 Hz. Participants were asked to move as little as possible and keep their gaze fixed on the computer screen in front of them, which displayed a fixation cross and occasional behavioral questions (see the Experiments section).

#### Preprocessing

The EEG recordings were analyzed using Matlab R2019b (The MathWorks). First, the following preprocessing steps were conducted using Letswave 6 (http://letswave.org) (Mouraux and Iannetti, 2008): high-pass filtering above 0.5 Hz with a 4^th^ order zero-phase Butterworth filter, 50 Hz bandpass notch filtering, down-sampling to 500 Hz, segmentation of trials from −1 to +1.5 seconds relative to stimulus onsets, baseline mean correction, and rejection of stereotyped artifacts using an Independent Component Analysis (ICA) decomposition (Bell and Sejnowski, 1995). Then, using Matlab, epochs were low-pass filtered below 30 Hz and trials with amplitudes reaching 80 *μ*V were rejected, leading to keep 491±17.3 and 490.2±16.27 (grand mean ±standard deviation) stimuli of intensities *I*_1_ and *I*_2_.

#### Linear regressions

We sought to determine if and how the Vertex Potential (VP) reflects the behavioral outcomes observed during TSL. The model which best approximated the participants’ behavior was considered (Bayesian model learning the TPs with a time constant *ω* = 8), and the VP was regressed on its key inferential outcomes. Two regressors were included in the analysis: the prediction error (see (8), known to affect sensory responses (Maheu et al., 2019;Strube et al., 2021)) and the confidence in the estimates, which weights learning in a Bayesian framework (Meyniel and Dehaene, 2017) (see (7)).

To ensure that the effects of confidence on EEG signals were not driven by confounding factors related to the prediction itself (*p*(*I*_1_|*y*_1:*n*_, *M_i_*, *ω_i_*) :≔ *p_n_*) (Meyniel, 2020), we first computed the *residual confidence* 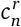 from the confidence *c_n_* by regressing out the predicted probability, its logarithm and its square as:

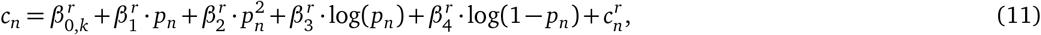

where *k* denotes the testing block index, *n* the trial index and *β^r^* the regression coefficients. The first coefficient 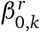 is a fixed intercept grouped by testing condition *k* (i.e. generative probabilities of the sequences). Then, for each participant, at each channel and at each time point from −0.5 to 1 second around stimulus onset, the EEG signal *z_n_* was regressed on the Bayesian prediction error (BPE) *e_n_* and residual confidence 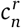 (omitting the dependence of the regressors upon the model *M_i_* and its parameter *ω_i_* for clarity):

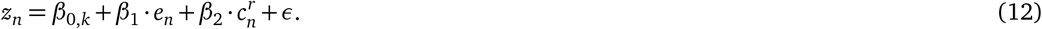

The regressions were computed across all available trials.

The two considered regressors – BPE and residual confidence – deduced from the optimal inference were not linearly related, enabling to compute and safely interpret the regression coefficients. To confirm that they are not collinear, we computed the Variance Inflation Factors (VIFs) for (residual) confidence against BPE (Sheather, 2009): 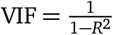, where *R*^2^ is the coefficient of determination obtained when linearly regressing (residual) confidence on BPE. Unless stated otherwise, ‘residual’ is assumed when mentioning confidence in this work. Significance of the regression coefficients across participants was assessed using one-sample *t*-tests against 0. Significance level was set to 0.05 and corrected for multiple comparisons across time points and selected channels (C3, Cz, FCz, CPz, C4) with the False Discovery Rate (FDR) correction.

## Supplemental information

Supplemental Information includes 4 figures and can be found with this article online.

## Acknowledgments

This work was supported by a Medical Research Council Career Development Award to FM (MR/T010614/1) and Wellcome Trust grants to BS. DM is a Research Fellow of the Fonds de la Recherche Scientifique - FNRS.

## Author contributions

DM, BS, AM and FM designed the experiments. DM collected and analyzed the data. DM and FM wrote the original paper draft, which was reviewed and edited by all co-authors.

## Data and code availability

The behavioral and EEG data sets will be made publicly available on an OSF repository upon acceptance and are also available from the corresponding authors upon request. The codes used to generate the model outcomes and analyze the data are available on GitHub (upon acceptance).

---

**Figure S1.**
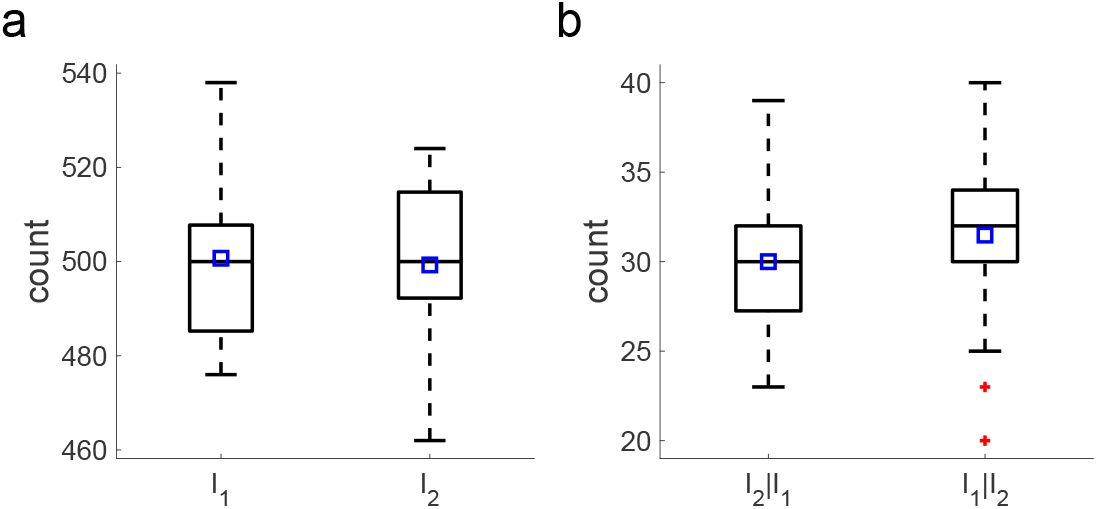
The number of stimuli from both intensities and the number of rated transitions are balanced. **a**, Numbers of stimuli from each intensity delivered to the participants along all the testing blocks. Participants received balanced numbers of stimuli from both intensities (mean difference between the numbers of *I*_1_ and *I*_2_: −1.419, *t*_30_ = −0.25, *p* = 0.804, Cohen’s *d* = −0.045). Blue squares: mean number of stimuli. **b**, Likewise, participants rated, on average, similar numbers of both types of transitions (mean difference of numbers of rated transitions from *I*_1_ and from *I*_2_: 1.484, *t*_30_ =1.054, *p* = 0.3, Cohen’s *d* < 10^−5^).

**Figure S2.**
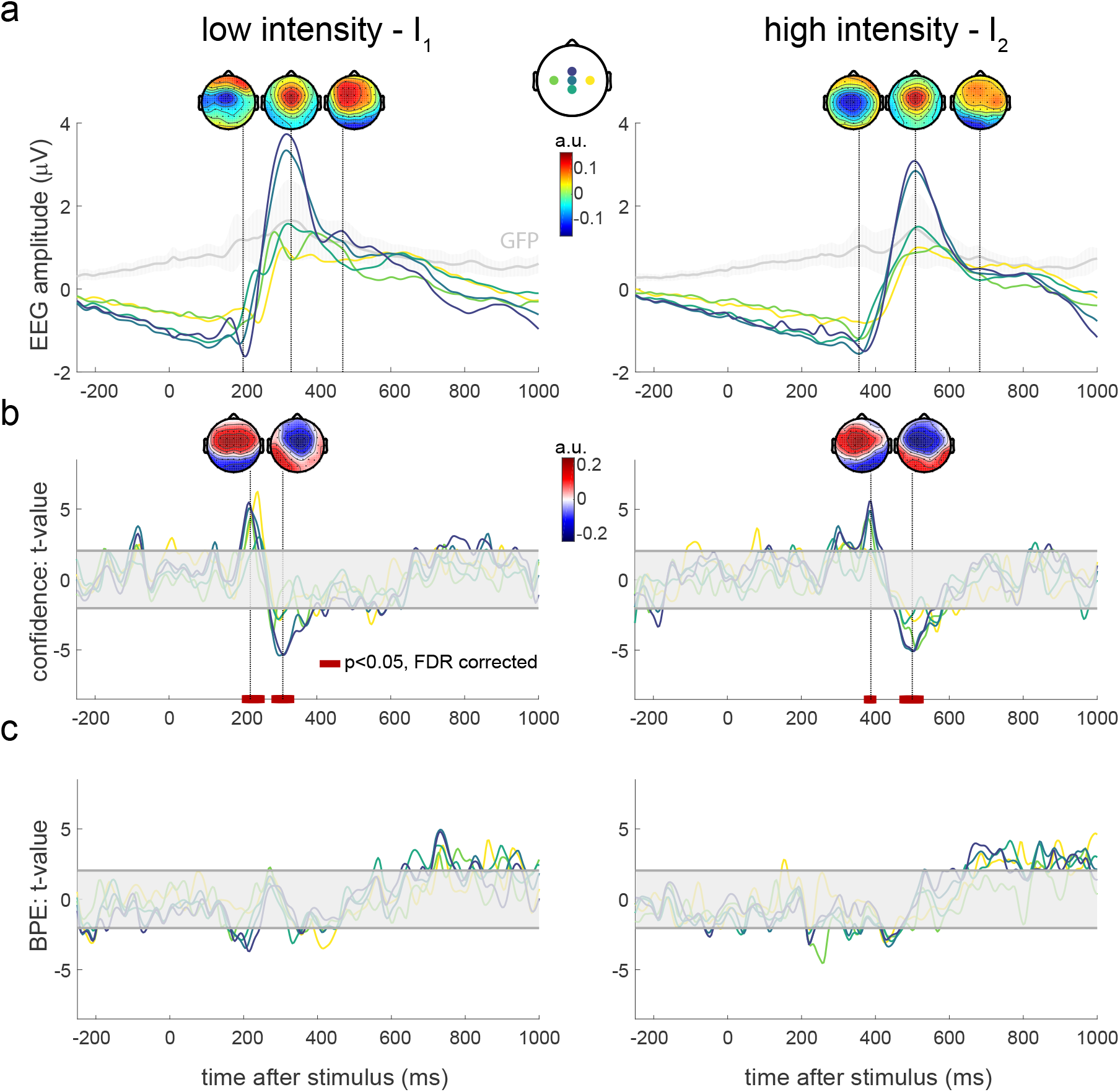
Links between the EEG responses and the Bayesian prediction error (BPE) and raw confidence. Similar to Fig. 5 but using the raw instead of residual confidence as regressor. **a**, EEG responses averaged over trials and blocks, for low (left) and high (right) stimulation intensities. Global Field Power (GFP) time courses are shown in gray, with shaded s.d. across participants. Labels of depicted electrodes: C3, Cz, FCz, CPz, C4. **b**, Encoding of raw confidence in the EEG responses – *t*-statistics for the regression coefficients associated with the model confidence. **c**, Encoding of BPE in the EEG responses – *t*-statistics for the regression coefficients associated with the BPE. In **b** and **c**, confidence and BPE are obtained from the model which best explains the participants’ behavior: a Bayesian model learning the TPs with an integration time constant of 8 stimuli. The shaded horizontal areas centered around 0 indicate the non-significant regions for *p* < 0.05, two-tailed. Red bars at the bottom of the plots show intervals where the regression coefficients are significantly different from 0 after False Discovery Rate (FDR) correction of the significance levels. BPE and raw confidence are not collinear: the average variance inflation factors (VIFs) of raw confidence against BPE =1.1 and 1.08 for *I*_1_ and *I*_2_ respectively, far below 5 (Sheather, 2009) (*R*^2^ = 8.92 and 7.36% when we regress the raw confidence on BPE).

**Figure S3.**
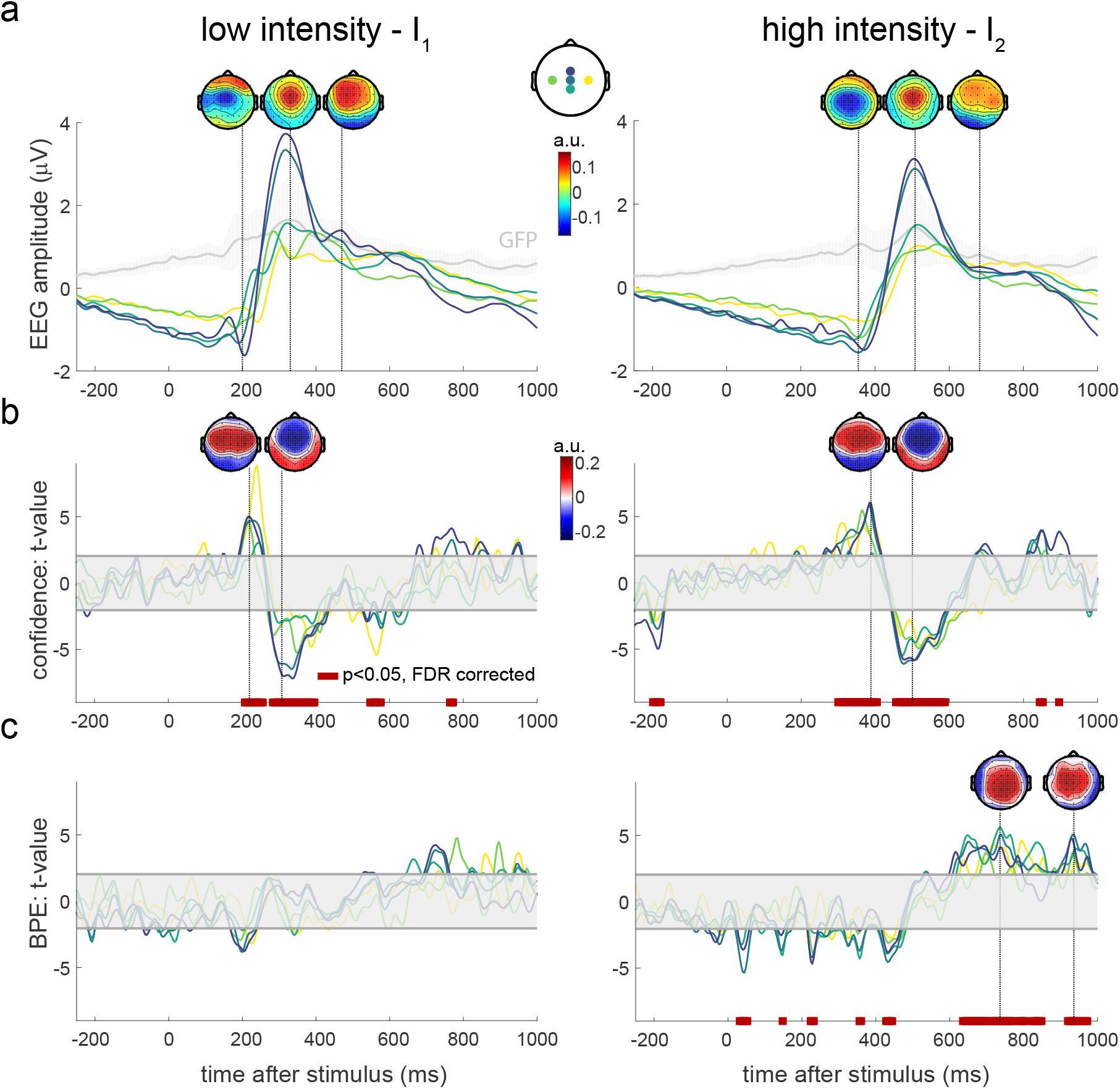
Links between the EEG responses and the Bayesian prediction error (BPE) and residual confidence. Similar to Fig. 5 but using a Bayesian model learning the IF instead of TPs. **a**, EEG responses averaged over trials and blocks, for low (left) and high (right) stimulation intensities. Global Field Power (GFP) time courses are shown in gray, with shaded s.d. across participants. Labels of depicted electrodes: C3, Cz, FCz, CPz, C4. **b**, Encoding of residual confidence in the EEG responses – *t*-statistics for the regression coefficients associated with the model confidence. **c**, Encoding of BPE in the EEG responses – *t*-statistics for the regression coefficients associated with the BPE. In **b** and **c**, confidence and BPE are obtained from the second model which best explains the participants’ behavior: a Bayesian model learning the IF with an integration time constant of 8 stimuli. The shaded horizontal areas centered around 0 indicate the non-significant regions for *p* < 0.05, two-tailed. Red bars at the bottom of the plots show intervals where the regression coefficients are significantly different from 0 after False Discovery Rate (FDR) correction of the significance levels. Again, BPE and confidence are not collinear: the average VIFs of confidence against BPE =1.0048 and 1.0058 for *I*_1_ and *I*_2_ respectively, far below 5 (Sheather, 2009) (*R*^2^ = 0.47 and 0.58% when we regress confidence on BPE).

**Figure S4.**
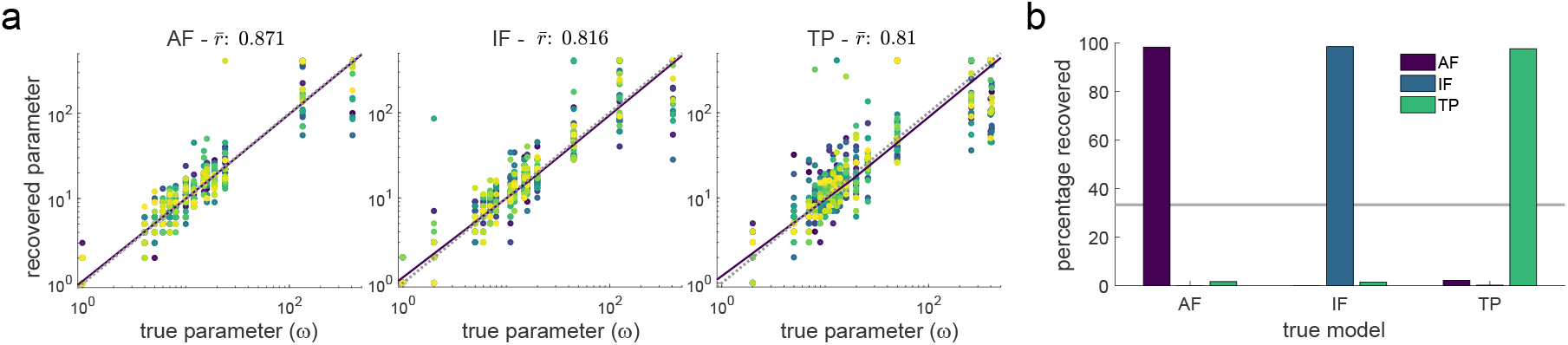
Parameter and model recovery for the Bayesian learner. To assess whether the three Bayesian models and their hyper-parameter (the integration time constant *ω*) are identifiable, we simulated data using each model with different parameters and fitted the models to these synthetic data like we did it to the behavioral reports. Since the model predictions (i.e. probability estimates) are deterministic for a given sequence, data were simulated by sampling probability estimates from the Beta distribution estimated at each time step (see Methods). For each model (learning AF, IF or TPs), we consider all the optimal time constants that were fitted to the individual behavioral data (Heald et al., 2021). Using each time constant and model, 30 synthetic data sets were built based on the same number of sequences and probability estimates as for the real participants (10 sequences were generated with the TPs indicated in Fig.1d and probability estimates were sampled every 15±3 stimuli). **a**, The parameter recovery analysis indicates that the integration time constant can be reliably recovered despite readout noise for all three models in our experiments (Pearson correlation coefficient between true and fitted *ω* =0.871, 0.816, 0.81). Scatter plots of fitted vs. true parameters, with one color per simulation (*n* = 30). Dotted line: identity, thick plain black line: linear fit. **b**, The model recovery shows that the three models are highly identifiable in our experimental setting, with 98.3, 98.6 and 97.6% of correctly recovered models for the model learning AF, IF and TPs respectively.

